# Phylogenomic analysis of SARS-CoV-2 genomes from western India reveals unique linked mutations

**DOI:** 10.1101/2020.07.30.228460

**Authors:** Dhiraj Paul, Kunal Jani, Janesh Kumar, Radha Chauhan, Vasudevan Seshadri, Girdhari Lal, Rajesh Karyakarte, Suvarna Joshi, Murlidhar Tambe, Sourav Sen, Santosh Karade, Kavita Bala Anand, Shelinder Pal Singh Shergill, Rajiv Mohan Gupta, Manoj Kumar Bhat, Arvind Sahu, Yogesh S Shouche

## Abstract

India has become the third worst-hit nation by the COVID-19 pandemic caused by the SARS-CoV-2 virus. Here, we investigated the molecular, phylogenomic, and evolutionary dynamics of SARS-CoV-2 in western India, the most affected region of the country. A total of 90 genomes were sequenced. Four nucleotide variants, namely C241T, C3037T, C14408T (Pro4715Leu), and A23403G (Asp614Gly), located at 5’UTR, Orf1a, Orf1b, and Spike protein regions of the genome, respectively, were predominant and ubiquitous (90%). Phylogenetic analysis of the genomes revealed four distinct clusters, formed owing to different variants. The major cluster (cluster 4) is distinguished by mutations C313T, C5700A, G28881A are unique patterns and observed in 45% of samples. We thus report a newly emerging pattern of linked mutations. The predominance of these linked mutations suggests that they are likely a part of the viral fitness landscape. A novel and distinct pattern of mutations in the viral strains of each of the districts was observed. The Satara district viral strains showed mutations primarily at the 3′ end of the genome, while Nashik district viral strains displayed mutations at the 5′ end of the genome. Characterization of Pune strains showed that a novel variant has overtaken the other strains. Examination of the frequency of three mutations i.e., C313T, C5700A, G28881A in symptomatic versus asymptomatic patients indicated an increased occurrence in symptomatic cases, which is more prominent in females. The age-wise specific pattern of mutation is observed. Mutations C18877T, G20326A, G24794T, G25563T, G26152T, and C26735T are found in more than 30% study samples in the age group of 10-25. Intriguingly, these mutations are not detected in the higher age range 61-80. These findings portray the prevalence of unique linked mutations in SARS-CoV-2 in western India and their prevalence in symptomatic patients.

**Importance:** Elucidation of the SARS-CoV-2 mutational landscape within a specific geographical location, and its relationship with age and symptoms, is essential to understand its local transmission dynamics and control. Here we present the first comprehensive study on genome and mutation pattern analysis of SARS-CoV-2 from the western part of India, the worst affected region by the pandemic. Our analysis revealed three unique linked mutations, which are prevalent in most of the sequences studied. These may serve as a molecular marker to track the spread of this viral variant to different places.

## Introduction

Severe acute respiratory syndrome coronavirus 2 (SARS-CoV-2) which is the causal agent for COVID-19, belongs to the category of betacoronaviruses. The respiratory illness caused by the virus varies from mild disease to severe disease and death (Guo et al., 2020; Tian et al., 2020). This virus has spread rapidly from Wuhan, China since late 2019, and on 11^th^ March 2020, WHO declared the disease caused by SARS-CoV-2 as a pandemic (Li et al., 2020). This crisis has rapidly escalated due to globalization and highly contagious nature of the virus. Till-date, more than 16 million people have been infected globally, and ∼0.6 million have succumbed to it (https://covid19.who.int/). Currently, after the USA and Brazil, India is in the third position on the basis of adversity of infection, and more than a million Indians have been infected until now. Among the various states of India, the Maharashtra state is a major hotspot for this disease, having around 1/5^th^ of total reported infections in India and thus needs more attention (https://www.covid19india.org/).

The genome sequences from across the world for SARS-CoV-2 are available through the Global Initiative on Sharing All Influenza Data (GISAID) platform since 12^th^ January 2020 (Shu and McCauley, 2017). The sequences of novel coronavirus (CoV) show a close similarity with severe acute respiratory syndrome-related coronaviruses (SARSr-CoV), and like SARS-CoV, it also utilizes ACE2 as their entry receptor (Zhang et al., 2020). The virus contains ∼30 kb positive-sense, single-stranded RNA genome, which encodes four structural and multiple non-structural proteins (Astuti 2020). Structural viral proteins form the capsid and the virion envelope. Non-structural proteins help in various stages of the virus life cycle like replication, followed by translation, packaging, and release (Lai and Cavanagh, 1997; Li 2016; Lu et al., 2020). Many mutations have emerged in the SARS-CoV-2 genome that can modulate viral replication, transmission and virulence efficiency (Jia et al., 2020; Pachetti et al., 2020). Genome sequencing of SARS-CoV-2, followed by the identification of genetic variants, have been reported from different parts of the world (Dorp et al., 2020). Recent studies have speculated that, under selective pressure, genetic variability accumulate and persist in the SARS-CoV-2 genome, for its better survival and transmission (Dorp et al., 2020). Most of the reported variants belong to Orf1a, Orf1b, S, N, and 5’ UTR region of its genome. Three sites in Orf1ab (region Nsp6, Nsp11, Nsp13), and one in the spike protein are characterized by an unusually abundant number of recurrent mutations that may indicate convergent type of evolution. Specific interest is imparted in the context of adaptation of SARS-CoV-2 in the human host (Dorp et al., 2020).

In this study, we explored the phylogenomic diversity among the SARS-CoV-2 from three different regions of Maharashtra, the western state of India. In particular, we generated whole-genome sequences from 90 samples and examined if there are any predominant and unique mutations responsible for different cluster formation among the present study samples. Further, we have also analyzed whether select variants are associated with the gender, age and symptoms.

## Materials and Methods

### Sample collection

Nasopharyngeal/throat swabs of suspected Covid-19 patients were collected in April and May 2020. Samples considered in the study confirmed positive by real-time PCR for SARS-CoV-2 at the National Center for Cell Science, B. J. Government Medical College Pune, or Armed Forces Medical College, Pune. The RT-PCR assay utilized WHO suggested primers and probes to target *E, ORF1b and RdRp* genes. Ethical clearance was taken from the Institutional ethical committee for the present study. Samples were anonymized except the information about gender, age, collection date, travel histories, and symptoms. The study sample details are given in Supplementary Data 1.

### RNA extraction and genome sequence

The COVID-19 patient samples that showed Ct value for E gene ranging from 18 to 24 employing RT-PCR were selected for genome sequencing. Viral RNA was extracted from 300 µl of viral transport media (VTM, Himedia, India) using QIAamp 96 Virus QIAcube HT Kit (Qiagen, Germany), following the manufacturer’s instructions. Extracted RNA present in 50 µl of elution buffer was immediately used for cDNA synthesis using random hexamer primers, following the manufacturer’s protocol (Qiagen, Germany). The extracted RNA and cDNA were stored at −80 °C until analysis. PCR based COVID-19 viral genome enrichment was performed using QIAseq SARS-CoV-2 Primer Panel Kit (Qiagen, Germany). A total of 98 primer pairs specific to COVID-19 were used for enrichment where primers pairs were distributed in two pools (Pool-1 and Pool-2). For multiplex PCR amplification, 5 µl cDNA of each sample was used as a template. The amplification was done with initial denaturation step at 98°C for 2 minutes, followed by 30 cycles of denaturation at 98°C for 20 sec, annealing/elongation at 72°C, and final extension step at 72°C for 20 minutes. Amplified PCR product of both the pooled primers set of the samples was run on 2% agarose electrophoresis gel. After EtBr staining, samples that showed amplification product with both the pooled primers were selected for further use.

The ∼400 bp amplicon product produced using both Pool-1 and Pool-2 from the same clinical sample was combined and purified using a 1X concentration of Agencourt AMPure XP beads (Beckman Coulter); the final product was eluted in 30 µl of elution buffer (Qiagen, Germany). The purified product was quantified using the HS DS DNA assay kit using Qubit (Invitrogen). For library preparation around 100-300 ng of purified amplicon were used. DNA library was made using QIAseq FX DNA Library Kit (Qiagen, Germany). To retrieve larger fragment library size, during digestion, 3 minutes incubation time was set followed by enzyme inactivation. Adapter ligation was performed using the supplied ligase enzyme by incubation for15 minutes at 20°C followed by enzyme inactivation by heating at 65°C for 20 minutes. After library preparation, library size selection and washing were carried out following manufactures ‘protocol using 0.8X and 1 X concentration of AmpureXP. The average library size was ∼500 bp. The purified library was sequenced by 2×250 bp chemistry using the Illumina MiSeq platform.

### Data analysis

Fastqc tool was used to check the quality of the raw paired-end sequences after sequencing (Andrews 2010). Adapter sequences and poor quality sequences were removed, and good quality sequences (Q>30) were selected using Trimmomatic (Bolger et al., 2014) for further analysis. Reference-based genome assembly was done using BWA (Burrows-Wheeler Aligner; Li 2013) to generate the consensus sequence. SARS-CoV-2 genome (NCBI GenBank accession MN908947.3) was used as a reference for mapping. Unmapped reads were discarded. Consensus sequences were used for checking completeness and coverage calculation. Cross verification of the assembly process using CLC genomics was achieved using the reference genome (Accession ID MN908947.3). Genome sequences were deposited in the GISAID database. The complete list of accession IDs of the study samples is listed in Supplementary Data 1.

### Phylogenomics analysis and functional evaluation of variants

Following the neighbor-joining method, a phylogenetic analysis was carried out to understand the relationship among the study sequences, with 1000 bootstrap. SARS-CoV-2 isolate Wuhan-Hu-1 (NCBI GenBank accession MN908947.3) was included as a reference genome in the tree. To identify the mutations and their position, a variant analysis was performed for all the samples utilizing SARS-CoV-2 sequence Wuhan-Hu-1 (NCBI GenBank accession MN908947.3) as a reference. The protein variants identified in the clade were assessed to know their functional effects using PROVEAN (Protein Variation Effect Analyzer) program, considering the protein sequences of the Wuhan-Hu-1 genome as reference and a default threshold value of −2.5 (Choi and Chan, 2015). It provided a universal method to calculate the functional effects of protein sequence variations that might be amino acid substitutions, deletion, and insertions at single or multiple levels. The pair-wise alignment-based score was calculated to estimate the sequence similarity change of the query sequence to a protein sequence homolog before and after the insertion/substitution of an amino acid variation to the query sequence.

### Structural and bioinformatics analysis of SARS-CoV-2 variants

Multiple sequence alignment: ClustalOmega (Sievers et al., 2011) and MUSCLE (Edgar 2004) as multiple sequence alignment tool were used to align protein specific regions. 3D structures of protein were retrieved from PDB database. Specifically, following PDB were used: Spike protein: 6MOJ, 6MIV, 6LXT, Orf3a: 6XDC; RdrP: 6×2G, 7C2K; and Nucleocapsid: 6WJI. Structural mapping and analysis of mutations was carried out in PYMOL (DeLano et al., 2002).

### A comparative study among the Indian samples

The phylodynamic analysis was done by Nextstrain pipeline following the standard protocol (Hadfield et al., 2018). The dataset of Indian samples till 24^th^ June 2020 was downloaded from the GISAID database and used for the analysis. A total of 943sequences from different regions of India were reported till that day. The details of samples used from the GISAID database, including sample id, sampling time, and submitting Institutes are listed in Supplementary Data 2. In addition, 90 samples from the present study were also included in the analysis. In this pipeline, all the sequences, including our study samples, were aligned using MAFFT (Multiple alignments using fast Fourier transform) (Katoh and Toh, 2008). Using IQTREE, the phylogenetic tree was made by the Augur tree implementation. Furthermore, the raw tree was processed with Augur for generating TimeTree, annotating ancestral traits, inferring mutations, and finally identifying clades (Nguyen et al., 2015). The resulting tree was viewed using Nextstrain. After phylodynamics and clade reclassification, only samples that were collected during April and May 2020 from other parts of India with a suitable representation were selected for comparative study.

### Statistical analysis

Relationship/association among the mutations with respect to their presence and absence in study samples was determined by Unweighted Pair Group Method with Arithmetic Mean analysis (UPGMA). The binomial and one way ANOVA was used to check the significance level of effect of gender, age, symptoms of the study samples.

## Results

### Demographics and data analysis

A total of 90 COVID 19 positive samples were sequenced, and were collected from Maharashtra, India, primarily from Pune, Satara, Nashik and Kolhapur districts. The age of the patients selected in the present study ranged from 2-78, with 80% patients were in the age range of 30-60 years. More than 1000x coverage and almost 99.99% genome completeness was achieved for all the study samples with respect to the reference genome, i.e., Wuhan COVID 19 genome. The metadata for the specimens have been deposited after sequencing in the public domain (Supplementary Data 1).

### Identification of phylogenomics cluster and responsible mutation pattern

A phylogenetic analysis of the viral genome sequences in the present study performed using Wuhan SARS-CoV-2 isolate’s genome (NCBI GenBank accession MN908947) as reference. The sequences analyzed in the study formed four clusters. Except for cluster 1, all other clusters were rooted from a single clade. Within this clade, clusters 2, 3, and 4 formed major groups (Figure 1). Cluster 1 formed by six sequences in the present study showed a close association with Wuhan reference sequence, indicating that these strains were genetically closer to the original sequences (Wuhan, China). Interestingly, ∼45% of the sequences in the study fell in cluster 4 which is most distant from cluster 1 (Figure 1). Cluster 2 and 3 formed the other two significant groups, each consisting of 18 sequences.

**Figure 1:**
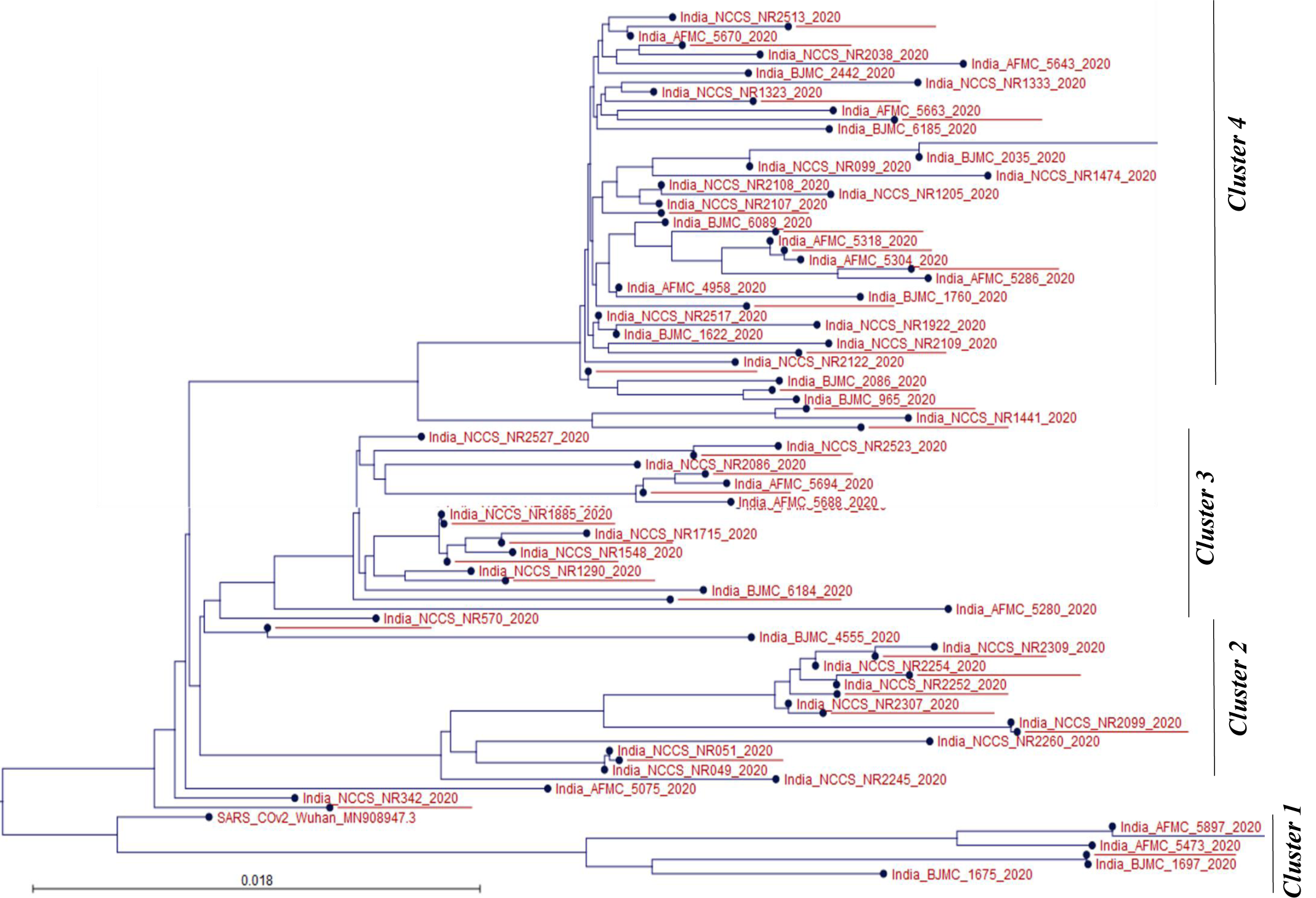
Phylogenetic analysis of the sequences used in the present study. Whole-genome sequences of the study sequences along with Wuhan SARS-CoV-2 genome (NCBI GenBank accession MN908947.3) as reference genome are used for the analysis.

To identify the basis of the formation of different phylogenetic clusters by the sequences of the present study, all the SNPs at different locations of the 90 genomes were mapped, followed by a heat map analysis based on the presence and absence of SNPs (Figure 2). Interestingly, a distinct pattern of mutations in different clusters was noticed in the present study (Figure 2). Of note, cluster 1 in the phylogenetic tree showed a distinct pattern of mutations compared to other clusters. Mutations C6312A, G11083T, C13730T, C23929T, and C28311T, were present in all the sequences of cluster 1, making them characteristic for this cluster 1 (Figure 2). The major cluster, i.e., cluster 4, was rooted from the same clade where clusters 2, 3, and 4 were also formed and variants C241T, C14408T, and A23403G were found to be key for the formation of this particular clade (Figure 2). These variants were detected in all the sequences, except six sequences of cluster 1. In addition to these variants, SNPs C313T, C5700A, G28881A were explicitly present in more than 95% sequences belonging to cluster 4, i.e., supercluster, and were variants for the formation of cluster 4 (Figure 2). This unique pattern of mutation, i.e., the simultaneous presence of the three mutations (C313T, C5700A, G28881A), has not been reported so far. Only mutation C313T and G28881A have been reported separately. Variants C3634T, C15324T were unique for all the sequences belonging to cluster 3, which were found in the region of Orf1a and Orf1b, respectively. Another major cluster, i.e., Cluster 2, where mutation C18877T, G25563T, C26735T, were unique and were found in ∼90% of the sequences of this cluster (Figure 2).

**Figure 2:**
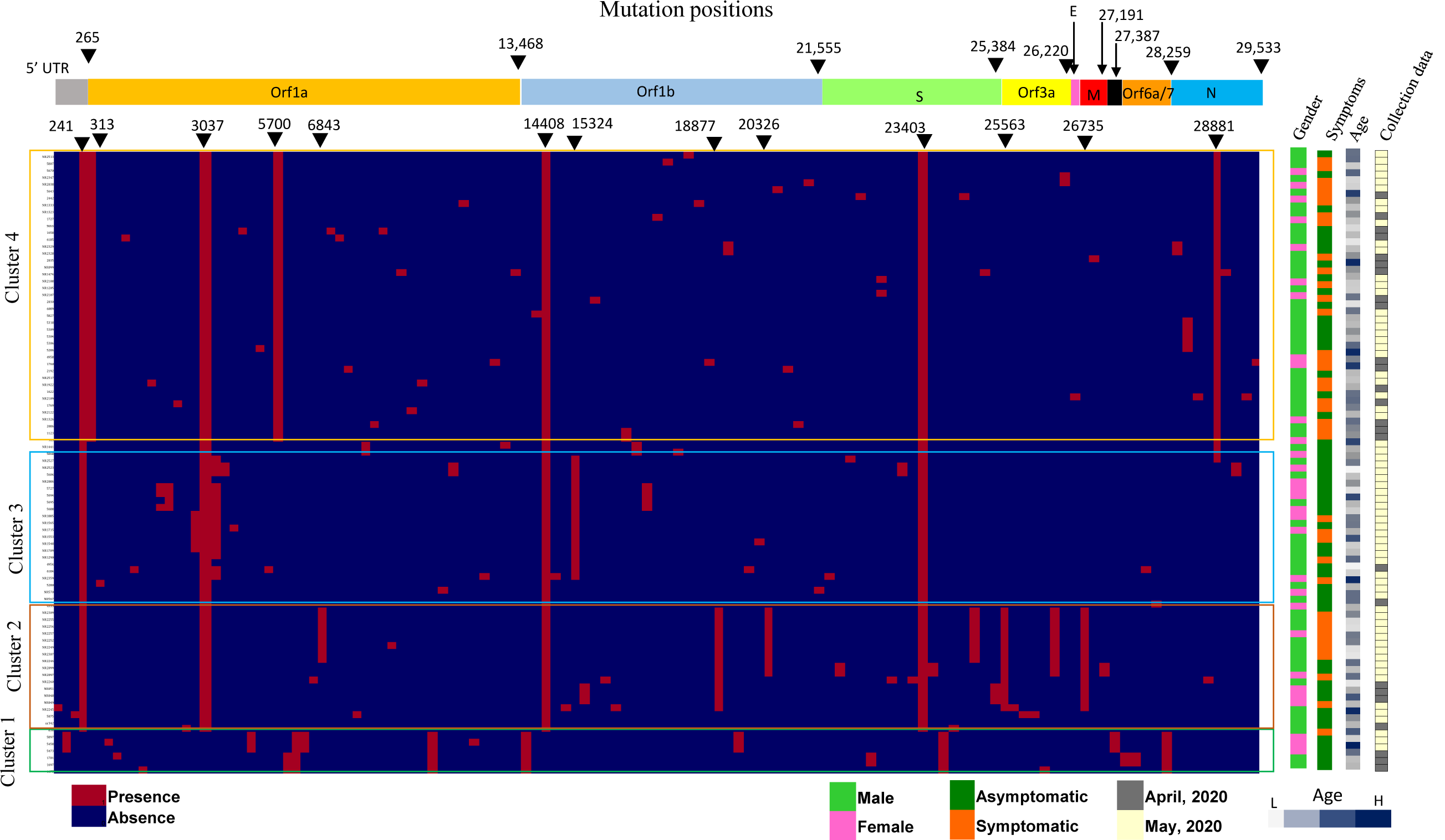
Heat map representing all the mutations found in the present study. Red color indicates the presence of the mutation, and blue color indicates the absence of mutation. The first column represents the mutation position in the nucleotide level, and the first row represents samples Id of the present study.

### Mutations and functional consequences

A total of 125 SNPs/variants were found in the present study. Most of the SNPs were found in Orf1a (46 SNPs) followed by Orf1b (31 SNPs), S (17 SNPs), N (10 SNPs), ORF3a (8 SNPs), 5′ UTR (4 SNPs), M (3 SNPs), E (1 SNP) regions of the genome (Supplementary Data 3). Among these variants, four were found in more than 90% study samples, i.e., C241T (5′ ÚTR), C3037T (ORF1a), C14408T (Pro4715Leu), and A23403G (Spike protein, Asp614Gly). Three variants C313T (ORF1a), C5700A (ORF1a, Arg to Lys), G28881A (N protein, Gly to Arg), were found in close to 50% of the sequenced samples. Mutations C3634T (Orf1a), C15324T (Orf1b), G18877T (NSP11 region of Orf1b), G20326A (Orf1b, Val6688Ile), G25563T (APA3_viroporin region ORF3a, Asn57His), G26152T (APA3_viroporin region ORF3a), C26735T were found in more than 20% of the study sequences (Supplementary Data 3). In addition, there were 100 other mutations that were found at a lower frequency (1-10%).

Protein Variant Effect Analyser (PROVEAN) was used to understand the functional consequences of non-synonymous variants formed due to amino acid substitution. A database for SARS-CoV-2 was constructed using the protein sequences of the WH1 reference genome (NC_045512) and the non-synonymous variants were analyzed. Two highly abundant (90%) variants of ORF1a/RdRP (Pro4715Leu) and Spike protein (Asp614Gly) were suggestive of neutral functional consequences as predicted by PROVEAN (Supplementary Data 3). Some mutations were found deleterious, based on PROVEAN analysis (Supplementary Data 3). However, the exact effect of this mutation should be validated experimentally.

### Region-wise mutation pattern

Region-wise mutation patterns among the viral sequences from Pune, Satara, and Nashik districts are depicted in Figure 3. Intriguingly, a specific pattern of mutation was found to be prevalent in all districts. Though mutations C241T, C3037T, C14408T (RdRp: Pro4715Leu), and A23403G (S protein: Asp614Gly) were dominant (80% of sequences) in all districts, Pune district sequences had additional predominant mutations C313T, C5700A, and GGG28881..28883AAC. Satara sequences, on the other hand, had unique mutations like G20326A (Nsp15: Val6688Ile), G24794T (S: Ala1078Ser), G26152T (Orf3a: Gly254Arg), and variants like C18877T and G25563T (Orf3a: Gln57His) in higher frequency, located towards the 3′ end of the viral genome, i.e., region NSP15 of Orf1b, S, and Orf3a. Sequences from Nashik district had unique mutations C190T, C1404T (Nsp1: Pro380Leu), T1872G (Nsp1: Phe536Cys), C3634T (Nsp3) and C5512T (Nsp3). It is interesting to note that unlike Satara sequences, the novel mutations observed in Nasik sequences were located at the 5′ end of the genome, i.e., in 5′ UTR, and Orf1a region of the genome (Figure 3).

**Figure 3:**
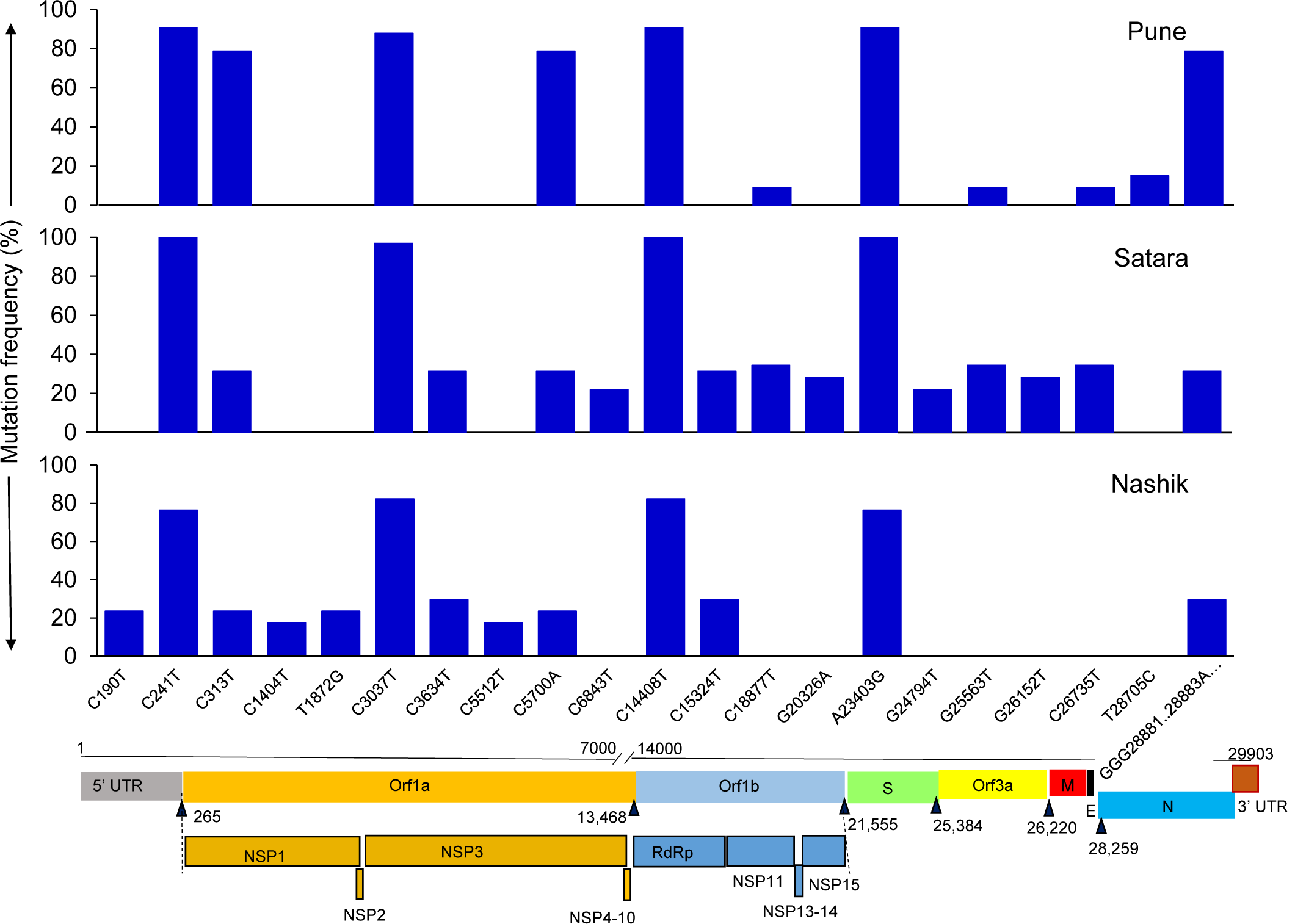
Region-wise mutation pattern of the Maharashtra state during the lockdown condition. The bottom-most bar represents nucleotide positions in various ORFs where mutations occurred on the genome (gene locations are based on Severe acute respiratory syndrome coronavirus 2 isolate Wuhan-Hu-1, NCBI GenBank accession MN908947.3 (https://www.ncbi.nlm.nih.gov/nuccore/MN908947)). Specific mutation observed in more than 5 samples are considered in this plot. Sample size: Pune (n=33), Satara (n=32), and Nashik (n=17)

### Mutation pattern with gender, age, and symptoms

The observed mutation pattern was further analyzed for any relationship with gender, age, and symptoms. With respect to gender, no predominant or unique mutation or mutation pattern was observed (Figure 4). A distinct pattern was observed in age-wise distribution (Figure 5). Its noteworthy that mutations C6843T, C18877T, G20326A, G24794T, G25563T, G26152T, and C26735T were not detected in the age range of 61-80 (Figure 5). Out of these specific seven mutations, C6843T, G20326A, G24794T, G25563T, and G26152T are non-synonymous mutations (Supplementary Data 3). These mutations are less prevalent (5-10%) in the age range of 26-60. However, in the age range of 10-25, the mutations were found significantly (p<0.05, Supplementary Data 4) at a higher proportion (>30%) (Figure 5). This indicated that though these mutations (i.e., C6843T, C18877T, G20326A, G24794T, G25563T, G26152T and C26735T) are observed in the age range 26-60, they are more prevalent in the age range of 10-25.

**Figure 4:**
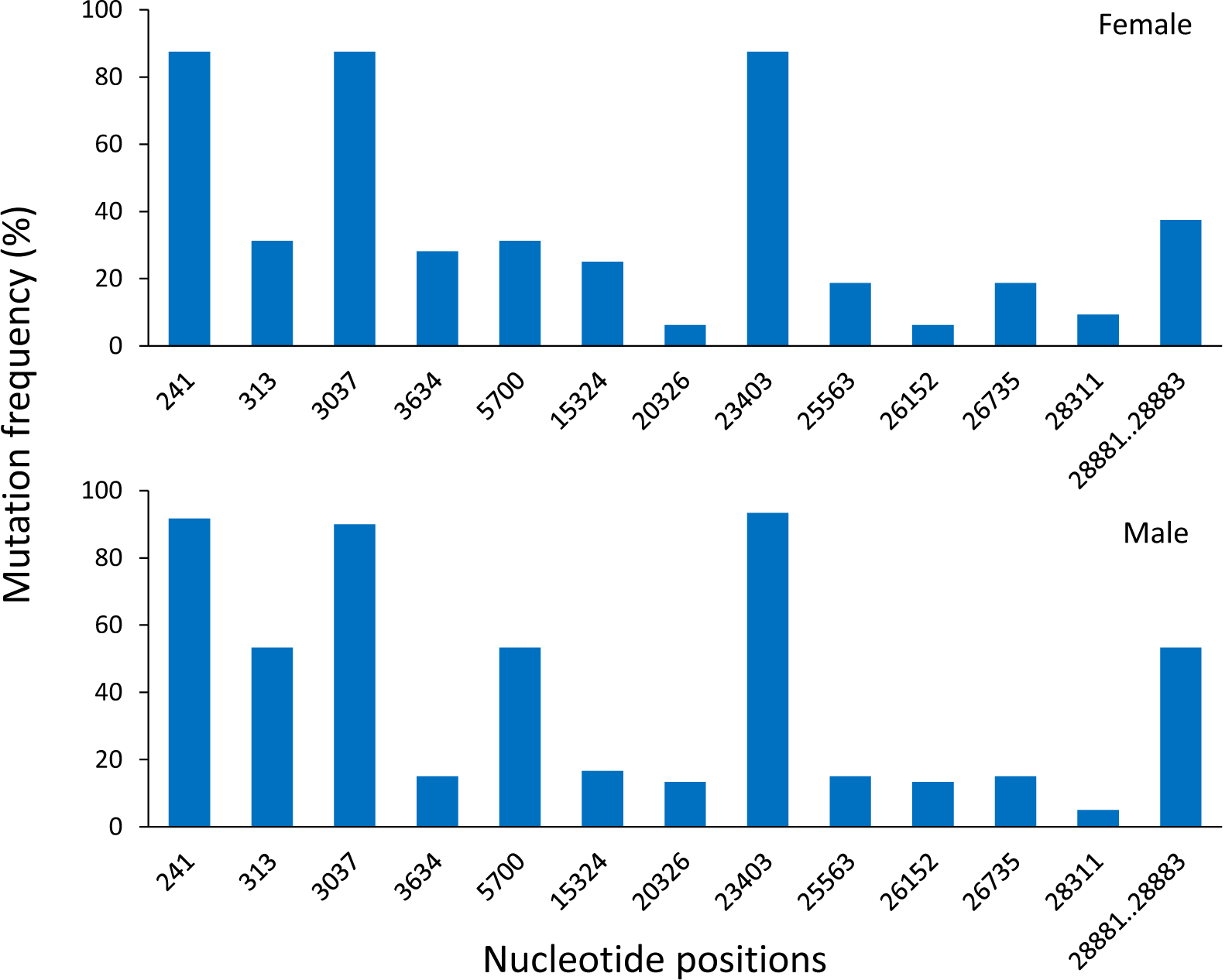
Gender-wise mutation pattern detection. The bottom part of the bar plot represents a nucleotide position where mutations occur on the genome. Specific mutation observed in more than 5 samples are considered in this plot. Sample size for female (n=31) and male (n=51).

**Figure 5:**
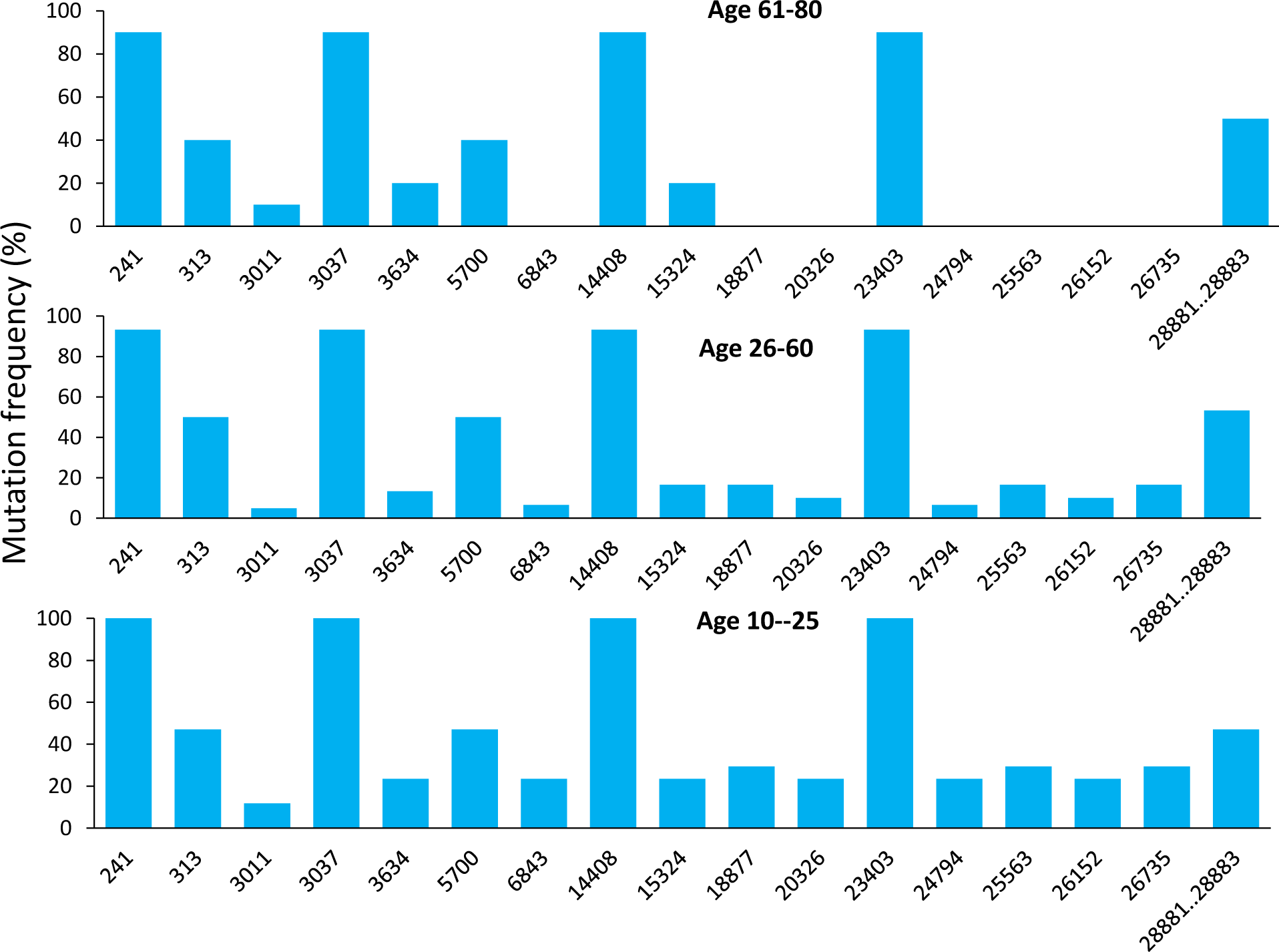
Age-wise mutation pattern identification. The bottom part of the bar plot represents a nucleotide position where mutations occur on the genome. Specific mutation observed in more than 5 samples are considered in this plot. Sample size for age range 10-25 (n=18), 26-60 (n=60) and 61-80 (n=10),

The relative changes in the frequency of mutations at positions C313T, C5700A, G28881A was observed when symptomatic and asymptomatic individuals were compared (Figure 6). The proportion of mutations C313T, C5700A, and G28881A were found relatively higher (∼80%) in symptomatic subjects as compared to asymptomatic (40-50%) (Figure 6). Interestingly, these differences in mutations between symptomatic and asymptomatic individuals were more pronounced females compared to males (Supplementary Figure 1 and 2). In the case of symptomatic females, the relative frequency of these mutations was close to 60% whereas for asymptomatic females, it was bout less than half, i.e.., 15-20%. In contrast, in case of males relative changes of these mutations, i.e., C313T, C5700A, and G28881A was observed less between symptomatic and asymptomatic individuals (Supplementary Figure 2).

**Figure 6:**
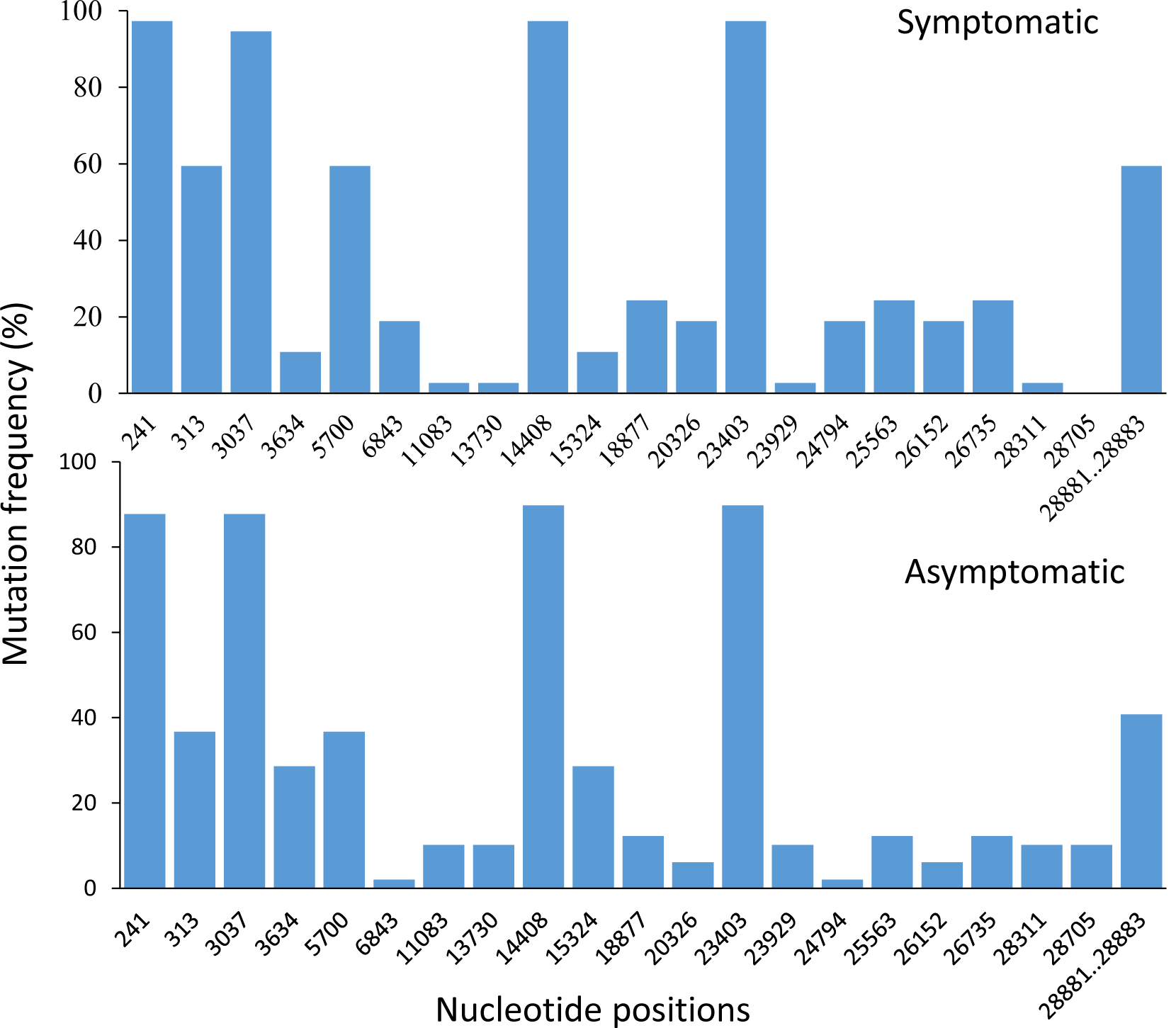
Symptom-wise mutations’ pattern identification. Specific mutation observed in more than 5 samples are considered in this plot. Sample size for patient of symptomatic (n=37) and asymptomatic (n=49).

### The pattern of mutation: Maharashtra compared to Indian scenario

Phylodynamic and clade reclassification of the previously sequenced genomes from India (943 genomes) and our study (90 genomes) was performed using Nextstrain. Notably 50% strains belonged to the 20A clade followed by 20B, 19A, and 19B (Supplementary Figure 3). The temporal changes in the appearance of different clades were further investigated. Data obtained from previous studies on six Indian states which were generated from the sample collected during April–May 2020 were only considered. The temporal change was studied by calculating proportions of sequences belonging to the five types in each of the two months under consideration. It was found that in capital Delhi (Northern part of India) during both the months, there were no changes. Only 19A type was dominant (100%), which was an ancestral China type (Figure 7). In Maharashtra (Western part of India), in April, 20B and 19A were prevalent where as in May, 19A proportion was reduced, and 20A and 20B were found in almost equal proportion. This indicated that the recent clade has become dominant with time in Maharashtra. Drastic changes were observed in Telangana (Southern part of India) (Figure 7). In April, only 19A clade was dominant (100%), but in month May, 19A type was found <5% and 20B and 20A were found to be more prevalent. Whereas the eastern part of India especially in Odisha, an opposite trend was observed in April, 20A type (90%) was dominating followed by 19A (10%) but in month May, 19A (50%) was prevalent followed by 20A, 20B (Figure 7). Thus, the overall pattern suggests that based on geographical location, the four sites of India showed unique SARS-CoV-2 prevalence. This may be one of the reasons for variable infection rates in different parts of this country.

**Figure 7:**
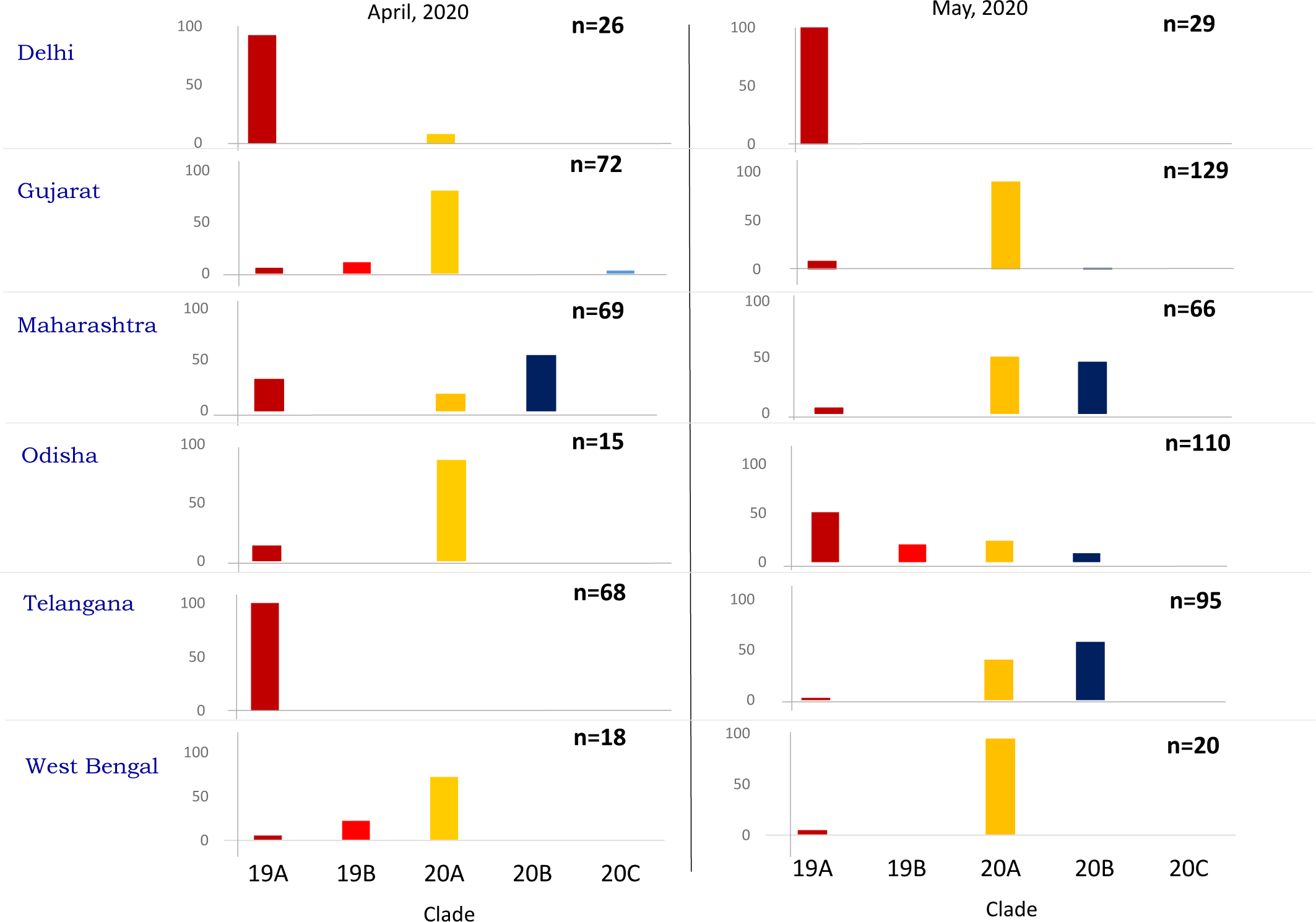
Temporal change in occurrences of five major types of SARS-CoV-2 (based on recent reclassification method) in different states of India. Sequences used for this analysis, retrieved from the GISEAD database.

## Discussion

In the present study, the goal was to explore the phylogenetic relationship among the Indian strains and to characterize the prevalent and unique genetic signature/pattern of the local strains sequenced from patients belonging to different regions of Maharashtra, the western part of India. A total of 90 viral whole-genomes were sequenced using nasopharyngeal and throat swabs. Recently, the pandemic caused by SARS-CoV-2 from Wuhan (China), clade O (Forster et al., 2020), has been reclassified as a 19A clade, which thereafter has evolved into multiple clades. Our phylogenetic analysis revealed 6 sequences with close proximity to the original Wuhan sequence, and others belong to three distinct clusters. This indicates that the sequences underwent several mutations in their genome, resulting in radiating phylogenetic clusters. In general, RNA viruses containing error-prone replication causes the mutations in their genetic material, which can be analyzed to track the viral evolution. The study until now proposes that SARS-CoV-2 arisen not very long before the first reported case of pneumonia in Wuhan (Wu et al., 2020). Having emerged recently, the observed level of diversity in SARS-CoV-2 is much lower compared to other known RNA viruses (e.g., dengue) where many subclones or lineages occur with multiple SNPs and associated functions (Wu et al., 2020). In the present study, 42 sequences belonged to supercluster 4 and mutations C313T, C5700A, G28881A were responsible for separating this supercluster from other clusters. Interestingly, all three mutations were found in the same genomes and UPGMA based cluster analysis (Supplementary Figure 4) indicating that they may be linked-mutations. Mutations C313T and G28881A have been reported previously in a separate study, but they were not present together with C5700A forming a supercluster. To the best of our knowledge, this finding in the present study is being reported for the first time.

About 50% of mutations in this study were synonymous, indicating null amino acid changes due to nucleotide substitution. At the same time, 50% of non-synonymous sites were detected, which might be due to convergent evolution. Of these non-synonymous mutations, the four most robust SNPs were present in above fifty percent frequency in the population, i.e., C5700A (>50%), C14408T (>90%), A23403G (>90%), G28881A (>50%) and are located in Orf1a, Orf1b, spike glycoprotein and Corona_nucleoca of N protein, respectively. Non-synonymous mutation at the A23403G position located in the Spike glycoprotein (Asp614Gly), which has a vital role in the binding of the virus to the ACE2 receptor in the host was observed.

The Asp614Gly mutation is very close to the Furin recognition site for cleavage of the Spike protein, which plays a decisive role in the virus entry (Korber et al., 2020; Hu et al., 2020). Interestingly, it has been shown that Gly614 mutant protein is more stable than Asp614 enabling increased transmission. Due to this, pseudovirions with Gly614 infected ACE2-expressing cells more efficiently than those with Asp614 (Zhang et al., 2020; Korber et al., 2020). We analyzed the Spike protein variants of Maharashtra stains using the known 3D structure of the spike protein (Wrapp et al., 2020; Wall et al., 2020). Asp614 site is located at the hinge region of the S1 and S2 domain and is shown to be involved in hydrogen bonding with T859. Any perturbation in this is proposed to destabilize the conformation of spike protein. Interestingly, this mutation was reported mostly in recent clades of SARS-CoV-2, which has a high frequency (more than 90%) in our study. A similar observation was made from other countries where the SARS-CoV-2 variant carrying the amino acid change in Spike protein at Asp614Gly position has become the most prevalent (Korber et al., 2020; Hu et al., 2020). This is in agreement with our results, which indicate similarity in the same pattern in three districts of Maharashtra. In addition to this prevalent mutation, the other 13 mutations in the Spike protein were also detected in the present study but at lower frequencies. Out of this G24794T, corresponding to Ala1078Ser lies in the S2 domain and located at the interface of trimeric oligomer and may affect the stability of the trimeric assembly of the spike protein. S2 domain is shown to be important for the fusion of viral membrane to the host-cell membrane. It was observed in 84 out of 90 tested samples (93.3%).This mutation is also reported in the European strains of SARS-CoV-2 (source: GISAID). We observed that this variant (6/84 samples) is coexisting together with Asp614Gly mutation in our data. Another SNP named C23929T, which is a synonymous, and neutral mutation was found in six sequences. Other SNPs were observed in one or two individuals/sequences only. Most of these mutations are unique for the Spike proteins and have not been reported before. Thus, it could be speculated that these new mutations might be emerging in our chosen geographical area. However, many factors may contribute, and their exact role in virus evolution or infection warrants to be explored further. Interestingly, no mutations in the furin cleavage site (PRRAR; 681-685) located at 23603-23617 (as per Wuhan-Hu-1 genome) which is responsible for the proteolytic cleavage of S protein and increased pathogenesis was observed in ourr samples.

In our sequences, a large number of SNPs (45 positions) were found in the Orf1a region of their genome, such as C313T, C3037T, C3634T, and C5700A. The C313T, C3037T, and C3634T are silent mutations, whereas a C5700A result in the Ala1812Asp substitution, and it is found in more than 50% of the sequences. The mutation C5700A in ORF1a results in missense mutation Ala1812Asp corresponding to Ala994Asp in the nsp3 protein was observed in 47.8% sequences. Interestingly, this mutation is reported in GISAID from other geographic locations of India. Orf1a containing of nsp 1 to nsp 10 plays an essential role in coping with cellular stress, retaining the functional integrity of the cellular components, and essential roles in the viral replication. Orf1a products play a vital role in viral structure and pathogenicity. Although the major mutations observed here were mostly silent, mutations in the Orf1 a region may affect viral structure and pathogenicity (Harcourt et al., 2004; Stobart et al., 2013). A number (35) of mutations were also found in the Orf1b region, among which C14408T, C15324T, C18877T, and G20326A were observed frequently. Mutation C14408T was found in more than 90% of the sequences, and it is a non-synonymous natural mutation resulting in Pro4715Leu substitution. This mutation has been dominant in the sequences across European countries. It is found in the RdRp region, which is involved in proofreading activities in the presence of other viral cofactors, like ExoN, nsp7, and nsp8 (Pachetti et al., 2020; O’Meara et al., 2020). It is anticipated that this mutation might affect proofreading capability. A minor alteration in the RdRp structure, without changing its catalytic activity, can also alter its binding efficiency with other cofactors such as ExoN, nsp7 or nsp8, and may contribute to change in the rate of mutation (Pachetti et al., 2020). Extensive studies are needed to understand the effect of mutations in RdRp, especially on viral replication. Two synonymous mutations at position C15324T and C18877T in NSP11 with moderate frequency (about 20%) were also observed in our study. We also observed another mutation with >6% frequency at position C13730T. Interestingly, this mutation is also reported in other states of India as well as in Singapore and Malaysia. This leads to a non-synonymous mutation Ala4489Val (Orf1ab) that corresponds to Ala97Val substitution in the RdRp protein. It has been suggested to substitute the alpha helix at positions 94-96 with beta-sheets, altering the tertiary conformation thereby might affect the fidelity of polymerase (Banerjee et al., 2020). It is also likely that these mutation lead to minor changes without affecting the functional activity of these proteins and thus adaptation in the host cell.

Putative cation channel encoded by Orf3a (about 274 amino acid long) is important for viral release, cell death, and inflammation. Our data indicated a significant number of mutations (G25552T, G25563T, C25886T, G26056T, G26065T, C26110T, and G26152T). All these mutations have been reported earlier in globally available sequences (source GISAID). It is known that the deletion of Orf3a in related SARS-CoV-1 leads to reduced viral titers and morbidity. Hence, it is important to understand how these Orf3a mutations contributing to COVID19 disease severity. Orf3a region translates a unique membrane protein with three-transmembrane, and G25563T variation in this region seems important (Cortey et al., 2020). We detected G255563T mutation at a frequency of 16.7% (n=15/90). This leads to mutation of Gln57 that lines the ion channel pore and forms the inner hydrophilic constriction to histidine. However, a recent study showed that the mutation Gln57His does not affect the assembly or functional properties of this putative cation channel (Kern et al., 2020).

SARS-CoV-2 N protein, a promising drug target for COVID-19, is a multifunctional RNA-binding protein required for RNA transcription and replication in virus. Thus play an important role in viral RNA genome packaging. Three distinct yet conserved domains of the N proteins are an N-terminal RNA-binding domain (NTD), a C-terminal dimerization domain (CTD), and intrinsically disordered central Ser/Arg (SR)-rich linker. Our study revealed a list of N protein mutations that is similarly reported in globally available sequences at GISAID. We noted that consecutive mutations at positions GGG28881--28883AAC on SARS-CoV-2 was observed in 47.8% sequences and is widely reported across the countries. These mutations lead to a non-frameshift substitution on Orf9. Notably, this has been reported in 23% of worldwide cases. G28881A and G28883C exist in the nucleocapsid gene and lead to variations in amino acid Arg203Lys and Gly204Arg, respectively. The Gly204Arg results in amino acid change with significantly different isoelectric point, and thus could potentially affect nucleocapsid protein structure and function (Pachetti et al., 2020). Non-synonymous mutation C28311T (Pro13Leu) in the nucleocapsid protein which is mandatory for the viral entry into the cells will likely have a significant effect on virulence was found in 6.7% of the sequenced samples.

The mutation C241T was found in 90% of all the sequences and is located in the 5′ UTR region and was found predominantly severely affected patients. It has also been shown to coincide with other mutations like C3037T (nsp3), C14408T (RdRP) and A23403G (S) in mildly affected, whereas in severe infections it coincided mostly with A23403G (S) (Biswas and Mudi, 2020). However, the role of this mutation has not been unraveled yet.

In the present study, a distinct novel pattern of mutations in closely located different districts of the same state was noted, perhaps due to strict, and effective lockdown. In Pune, a novel variant has overtaken the other strains. Satara and Nasik also have unique mutations but at a lower frequency. It would be interesting to study how these mutations affect viral fitness and virulence. Nonetheless, the lockdown resulted in an independent evolution of SARS-CoV-2 viral variants.

The host may also play a significant role in influencing the mutation pattern. One such host factor could be age, and age-wise specific pattern of mutations was detected in the present study. Mutations C6843T, C18877T, G20326A, G24794T, G25563T, G26152T, and C26735T were prevalent in more than 30% of the sequences were in the age group of 10-25 years. Surprisingly, these mutations were not found (0%) in a higher range of age, i.e., 61-80 years. The prevalence of viral variants might be dependent upon host-immunity and host genetic state, which is very different across elderly and young humans. Our results indicate that certain viral variants having specific set of mutations might be able to infect the young population more than the other less evolutionally advanced virus sub-clones. This is an exciting finding and demands further study in a larger cohort.

The mutations, i.e., C313T, C5700A, G28881A, were found more in samples from symptomatic subjects, specifically in symptomatic females. The same mutation was also found as a pattern in the present study supercluster 4. Hence, the linked-mutation, which is also evolving as a cluster in the present study, may have a role in exhibiting symptoms in a host-specific manner. Based on host immunity and physiology, it may be associated with the symptoms in that population where these mutations are prevalent. However, this observation needs to be validated with a much larger population-based study.

Additionally, in the Indian context, it was found that distinct sub-clones of virus were prevalent in different parts of India at the same time period. During April-May, 2020, the type 19A clade virus was predominant in the northern part of India (Delhi), which is an older clade whereas the western part (Maharashtra) was dominated by a more evolved clade, i.e., 20A, 20B. At the same time southern part (Telangana), where 19A clade was dominant in April, shifted completely to 20A and 20B in May 2020. Due to lockdown, most confounding factors (mobility of humans, inter-district transport, etc.) in the transmission of SARS-CoV-2 was restricted. Thus, we could assert that the prevalence of a specific viral variant in a region could be attributed to human host susceptibility for specific viral variants. In turn, this susceptibility seems to be based on mutations prevalent in the viral variants in that region. These factors also support the observation that indifferent parts of India, infection rate, viral load as well as mortality rate are variable during the same period.

## Acknowledgement

All the authors acknowledge the GISAID database and the concerned contributors of genomic data. The authors wish to thank the Department of Biotechnology, Ministry of Science and Technology, Government of India for the financial support and Dr. T. P. Lahane, Director, Department of Medical Education and Research for the permission to use the clinical specimens for this study. The authors would also like to appreciate the efforts of all those involved in the diagnosis, without their contribution it would not have been possible to generate the samples for sequencing.

## Contribution

Conceptualized and designed the study: YSS, AS, MKB, VS; Sample processing and sequencing: DP, KJ; Bioinformatics and statistical analysis: DP; Protein modeling: JK, RC; Data interpretation: DP, AS, YSS, JK, RC, VS, GL, MKB, RK, SS; Manuscript writing: DP with input YSS, AS; Manuscript improvement: AS, YSS, JK, MKB, VS, SK; Sample supply and coordination: RK, SJ, MT, SS, SK, KBA, SPSS, RM. All authors have read and approved the manuscript.

## Supplementary information

**Supplementary Figure 1:**
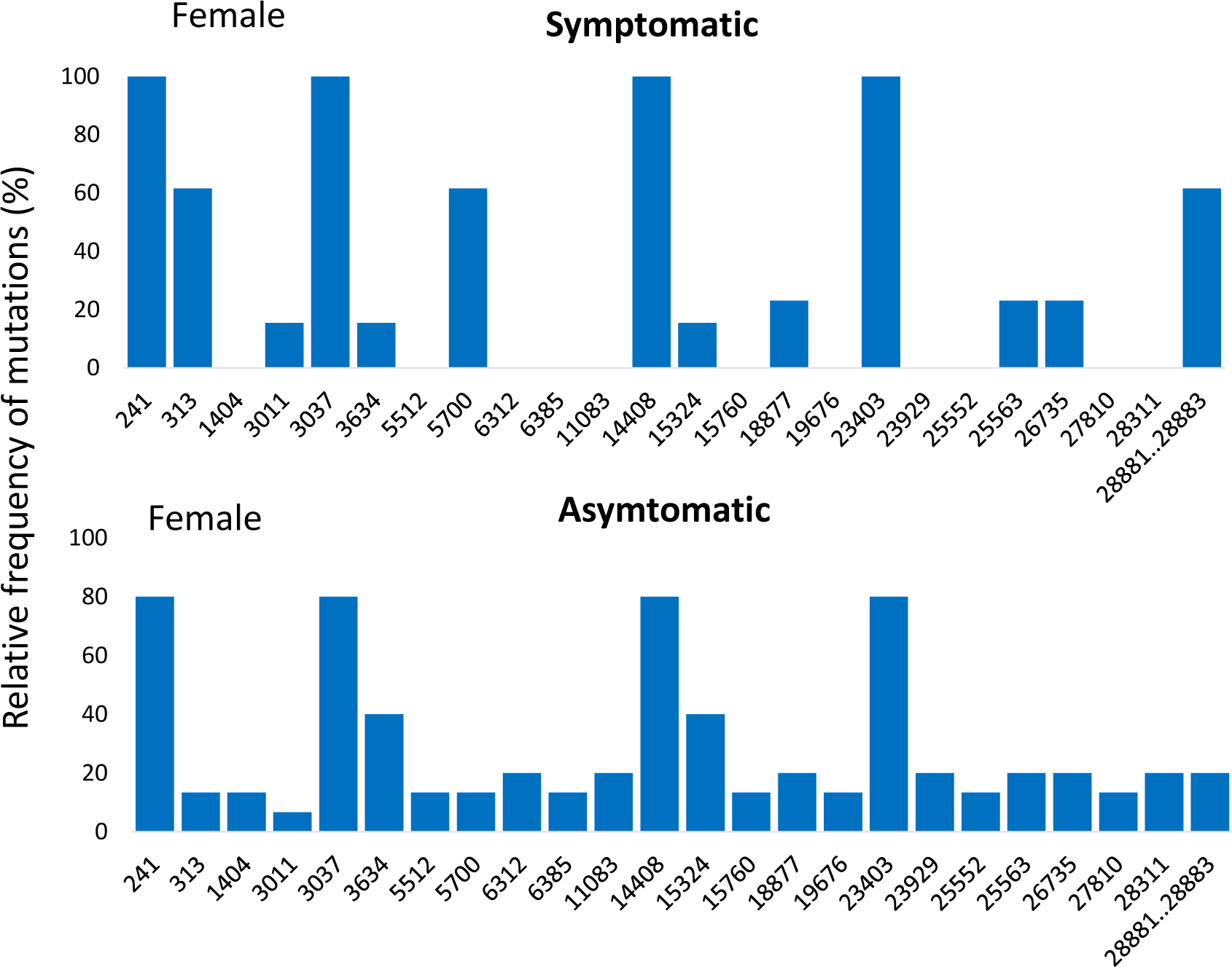
Symptom-wise mutation pattern identification females. Specific mutation observed in more than 5 samples are considered in this plot. Sample size for patient of symptomatic (n=13) and asymptomatic (n=15)

**Supplementary Figure 2:**
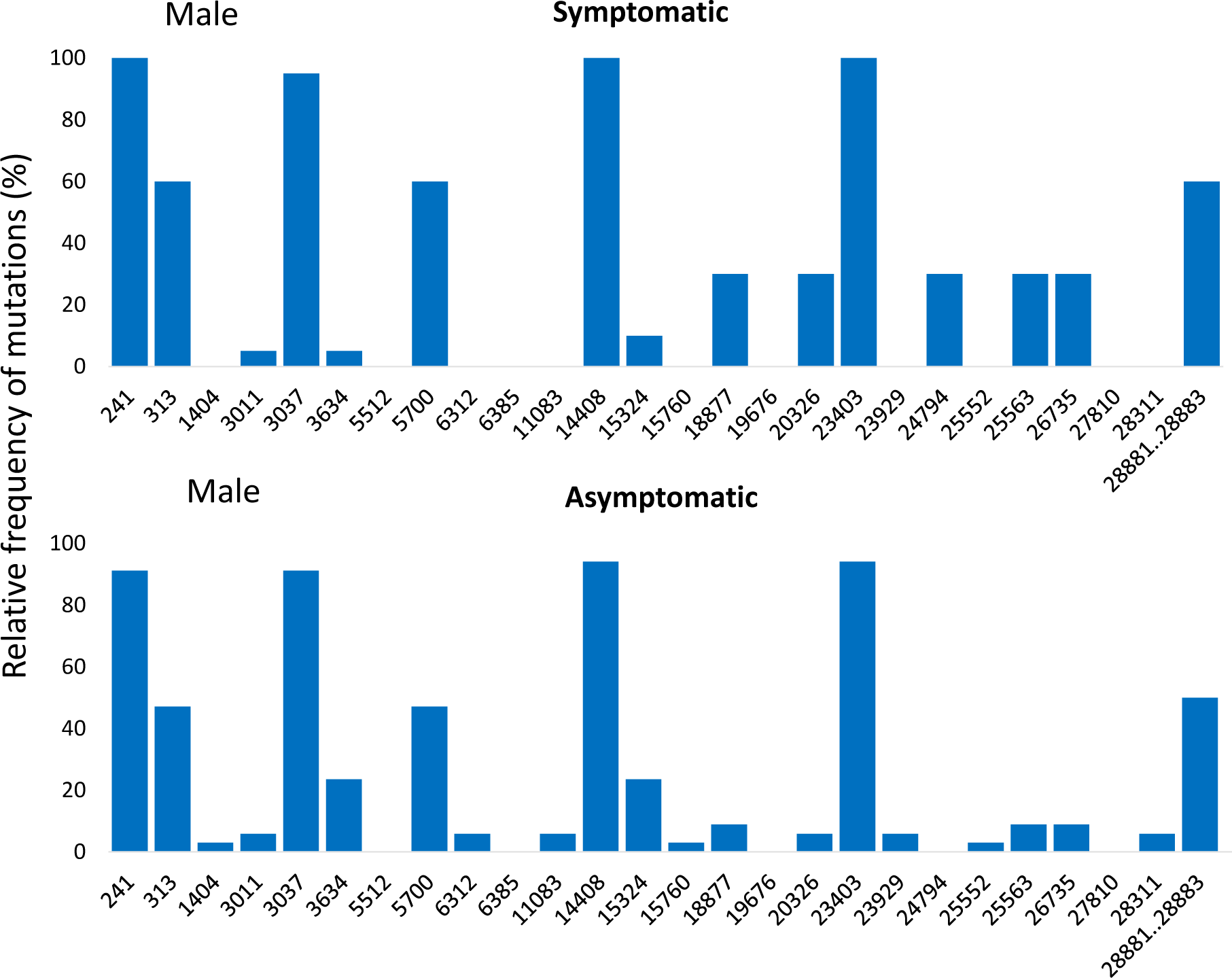
Symptom-wise mutation pattern identification in males. Specific mutation observed in more than 5 samples are considered in this plot. Sample size for patient of symptomatic (n=20) and asymptomatic (n=34)

**Supplementary Figure 3:**
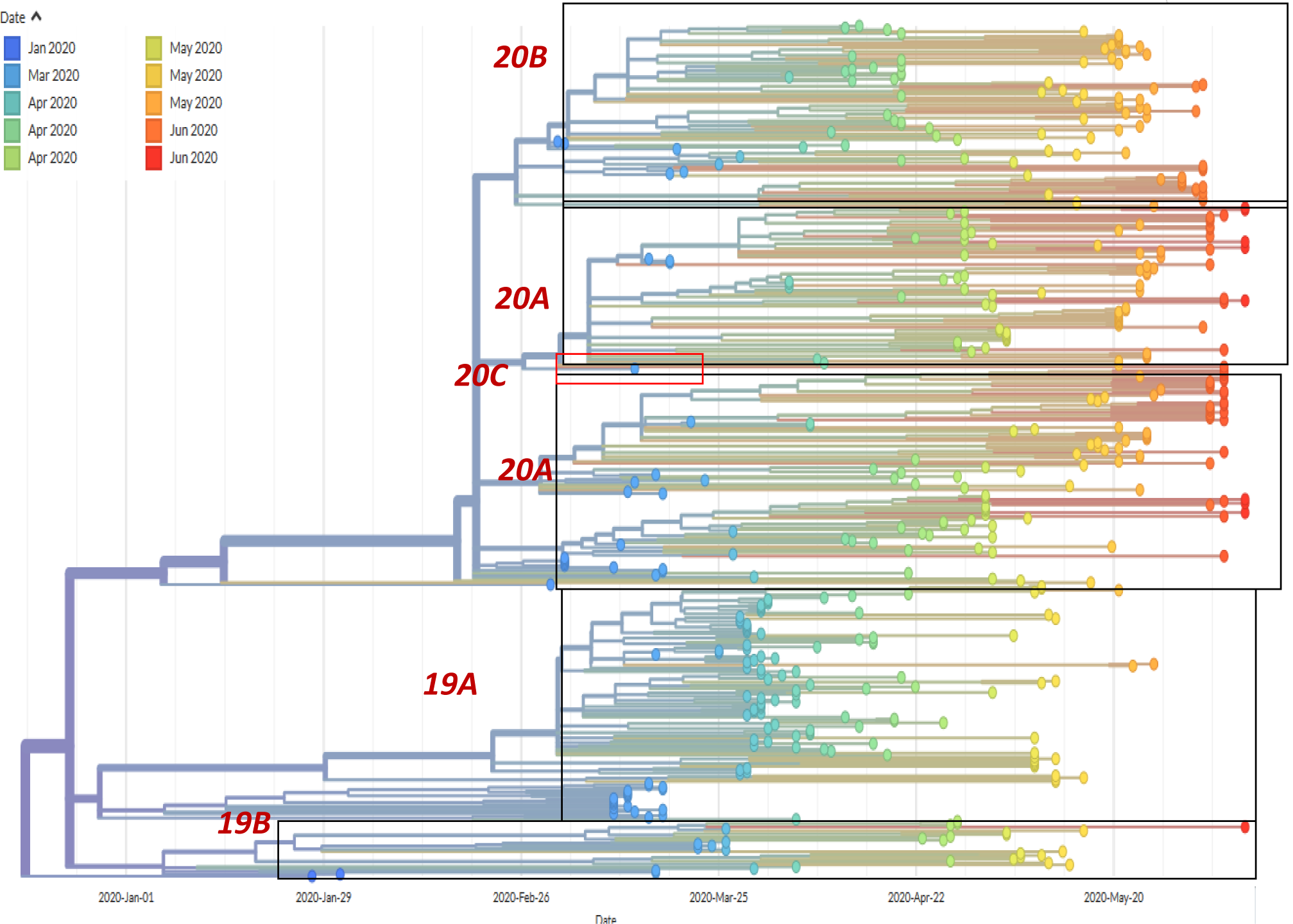
Phylodynamics clusters and the clades assigned based on the new reclassification method for the Indian SARS-CoV-2 Genomes (943 samples), including present study samples (90).

**Supplementary Figure 4:**
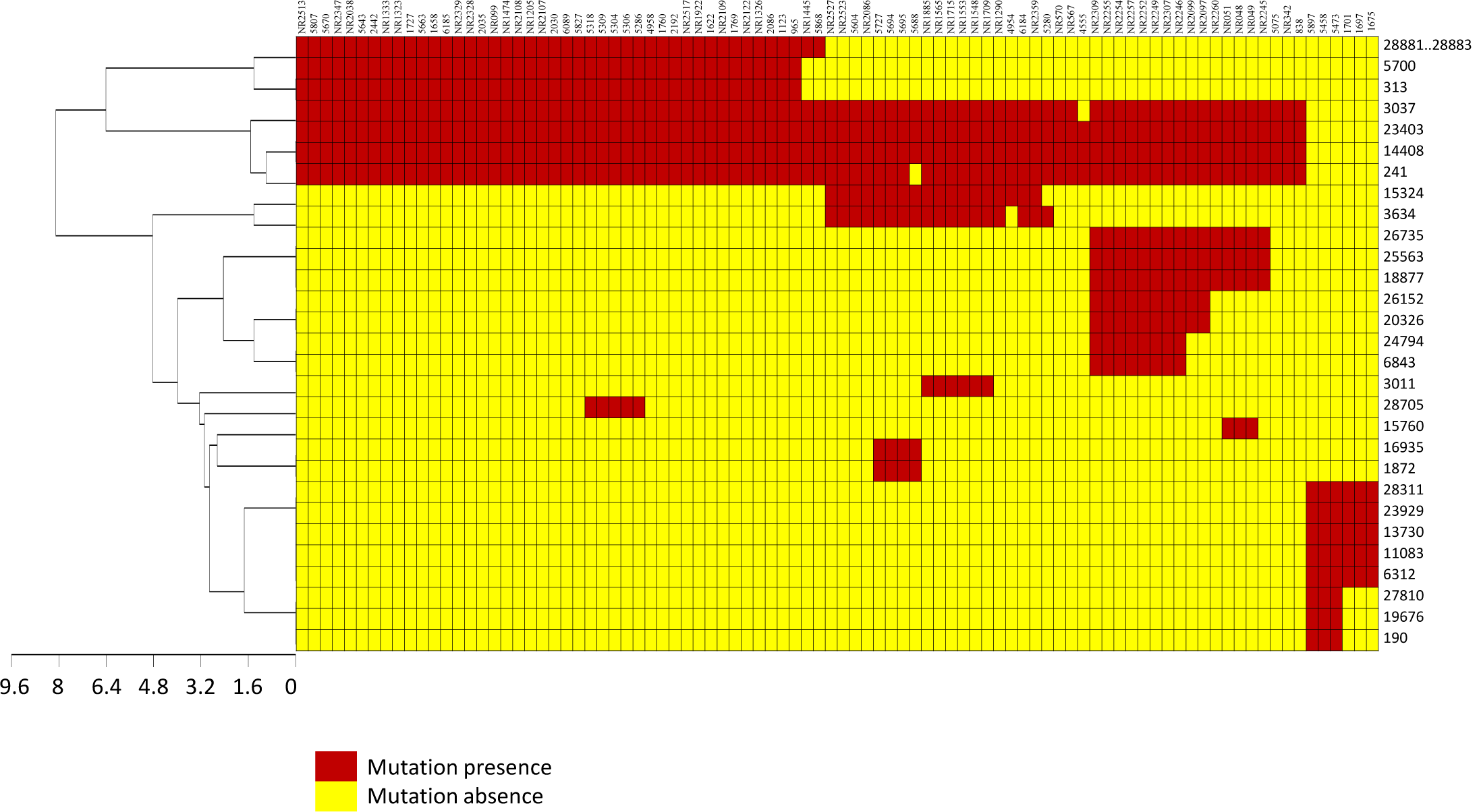
UPGMA-based cluster analysis for identification of closely associated mutation pattern

**Supplementary Data 1:** Metadata information of the present study samples

**Supplementary Data 2**: List of sequences deposited from India retrieved from GISAID database, which are included in the present study for comparative study and phylodynamics analysis

**Supplementary Data 3:** List of all mutations, abundance, and their mutational consequences

**Supplementary Data 4:** Statistical analysis

